# Clonally selected lines after CRISPR/Cas editing are not isogenic

**DOI:** 10.1101/2022.05.17.492193

**Authors:** Arijit Panda, Milovan Suvakov, Jessica Mariani, Kristen L. Drucker, Yohan Park, Yeongjun Jang, Thomas M. Kollmeyer, Gobinda Sarkar, Taejeong Bae, Jean J. Kim, Wan Hee Yoon, Robert B. Jenkins, Flora Vaccarino, Alexej Abyzov

**Author notes:** These authors contributed equally to this work (Arijit Panda, Milovan Suvakov, Jessica Mariani, Kristen L. Drucker, Yohan Park).

## Abstract

The CRISPR-Cas9 system has enabled researchers to precisely modify/edit the sequence of a genome. A typical editing experiment consists of two steps: (i) editing cultured cells; (ii) cell cloning and selection of clones with and without intended edit, presumed to be isogenic. The application of CRISPR-Cas9 system may result in off-target edits, while cloning will reveal culture-acquired mutations. We analyzed the extent of the former and the latter by whole genome sequencing in three experiments involving separate genomic loci and conducted by three independent laboratories. In all experiments we hardly found any off-target edits, while detecting hundreds to thousands of single nucleotide mutations unique to each clone after relatively short culture of 10-20 passages. Notably, clones also differed in copy number alterations that were several kb to several mb in size and represented the largest source of genomic divergence among clones. We suggest that screening of clones for mutations and copy number alterations acquired in culture is a necessary step to allow correct interpretation of DNA editing experiments. Furthermore, since culture associated mutations are inevitable, we propose that experiments involving derivation of clonal lines should compare a mix of multiple unedited lines and a mix of multiple edited lines.

## 1. Introduction

The CRISPR-Cas9 system has enabled researchers to precisely modify the sequence of a genome. The idea is borrowed from the defense mechanism of bacteria fighting against a virus attack (Jinek et al., 2012). The system uses an RNA-protein complex consisting of two main components: a Cas9 effector protein and a single guide RNA (sgRNA) targeting a sequence of about 20 nucleotides. Once the RNA-protein complex enters the cell, the sgRNA recognizes a complementary target sequence in the nuclear DNA and Cas9 makes double stranded break (DSB) in the DNA. The repair of the break may result in the sequence variation at the target site, such as an indel creating a loss of function mutation in a particular locus. If provided, a template DNA can be incorporated into the targeted site, to insert a desired variation or sequence (such as a tags or a selectable marker) in a particular position of the genome. Through this process, any DNA sequence can be precisely edited with an efficiency far superior to that of other endogenous genome editing methods like ZINC finger (Kim et al., 1996) or TALEN nucleases (Gaj et al., 2013). Currently, CRISPR-Cas9 system is used extensively to study the effect of a genetic variation within the human genome (Ran et al., 2013). Modified CRISPR-Cas9 systems can also be applied to induce single strand breaks, activate gene expression, and activate or repress non-coding elements (Hanna and Doench, 2020; Ma et al., 2015).

In a typical genome editing experiment, cultured cells are transfected with plasmids or transduced with a viral library (or libraries) encoding sgRNAs and the Cas9 nuclease. From the bulk cell culture, clonal subpopulations (e.g., clones with and without the intended edits) are isolated and expanded for further study. Of particular interest are studies that utilize human induced pluripotent stem cell (iPSC) lines. In this case, the lines are derived from somatic cells of an individual, and the CRISPR-Cas9 system is often used to study the phenotypic effect of a particular genome edit (such as a deletion, substitution, or an indel creating a loss of function mutation in a particular locus) on that individual’s genetic background. Typically, cells of the edited iPSC line are cloned, and clones with or without a given edit are compared and referred to as “isogenic” except for the induced mutation. This approach is often highlighted as particularly rigorous and powerful, in that it allows comparing the effect of single mutations in an otherwise unform genetic background.

However, the genomes of selected clones can be different. Differences can be introduced because of unintended edits of the Cas9 enzyme, so-called off-target effects (Boyle et al., 2017; Fu et al., 2013; Pattanayak et al., 2013). More importantly, genomic differences can arise from the natural accumulation of mutations in cells of the parental (iPSC) line. While these mutations are present in a small percentage of cells in a line, they can be disregarded as a potential source of phenotypic variation. However, upon cloning a cell from the lines those mutations will be manifested in all cells of each clonal derivative, becoming a potential source of phenotypic variation between clones. Off-target effects have been studied in multiple reports (Kim et al., 2015; Tsai et al., 2017) but less emphasis has been given to understand the effect of clonal selection and manifestation of preexisting somatic mutations in the clones (Assou et al., 2018).

Here we studied the effect of clonal selection in CRISPR-Cas9 editing experiments by analyzing data from three independent studies that were done in three different institutions (Yale University, Mayo Clinic, and OMRF/BCM). In each experiment, multiple clones were expanded, and single nucleotide variations (SNV) and copy number alterations (CNA) in each of the clones were analyzed.

## 2. Results

### 2.1. Experimental designs

Three independent experiments were conducted in three different institutions (**Fig. 1A**). In the experiment performed at Yale University, an induced pluripotent stem cell (iPSC) line, i3, generated from skin fibroblasts of an adult male, S1123-01, was subjected to editing. A locus in the *FOXG1* gene was targeted to produce a truncated, loss of function allele of the *FOXG1* gene. A double-stand DNA break in the *FOXG1* gene was generated by Cas9 using a gRNA binding to the sequence corresponding to N-terminal of the produced protein. When repaired by non-homologous end joining (NHEJ), an error prone DSB repair pathway, the sequence at the break may contain small deletion or insertions (indels). Post-editing, multiple clones were derived, expanded and, upon checking the target site with Sanger sequencing, one control (B10_CTR) and two clones with 1-bp insertion (A4 and A8) were picked for whole genome sequencing (WGS).

**Figure 1:**
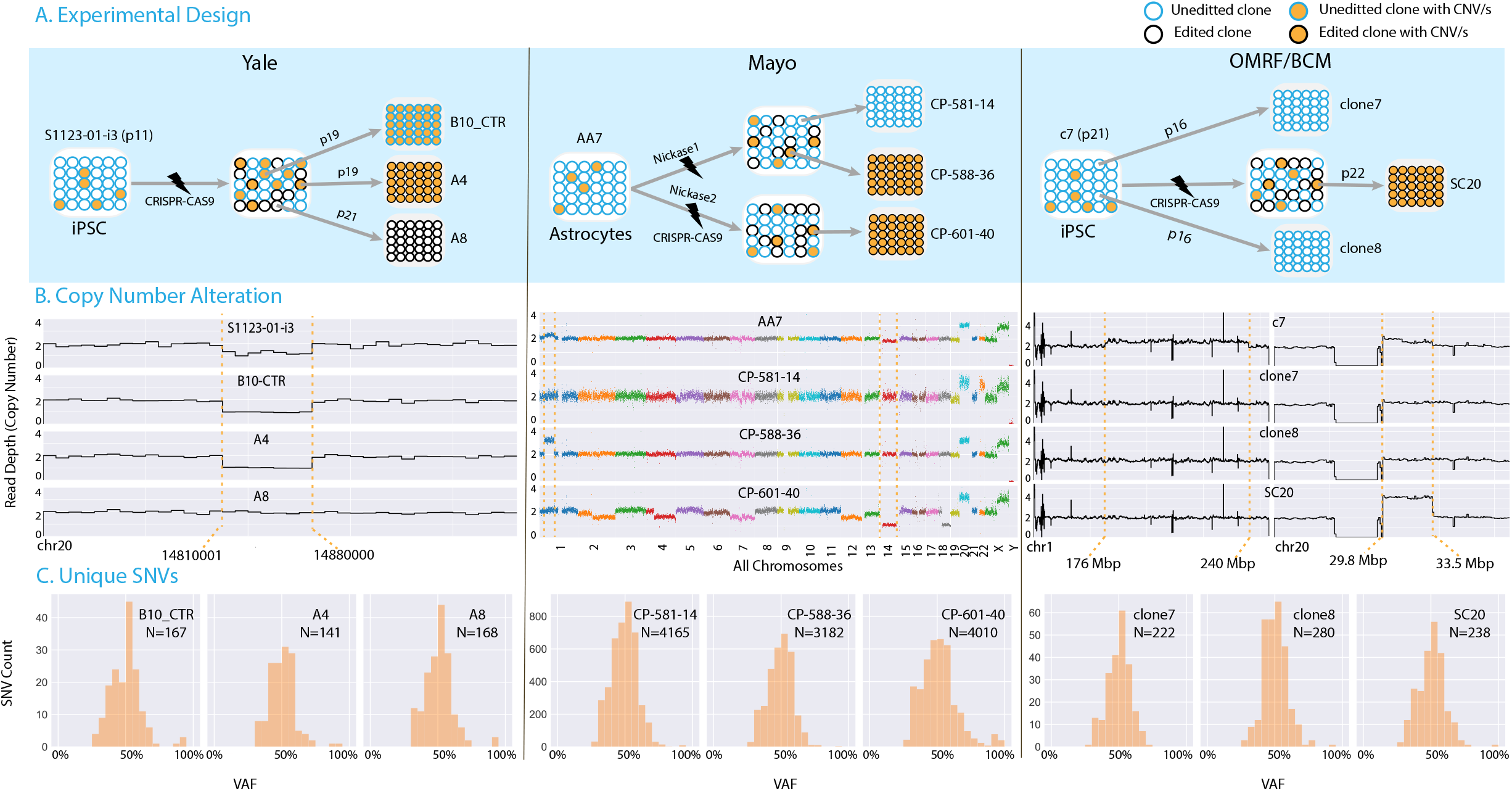
CRISPR-Cas9 edited clones are not isogenic. **A)** In three different experiments conducted by three different groups, CRISPR-Cas9 unedited (in blue circles) and edited (in black circles) single cell clones were analyzed. Orange filled circles represent clones with CNV. **B)** Copy number profiles for the analyzed samples. CNA inherited from unedited parental lines are shown between red broken lines. In the experiment conducted at Yale, the heterozygous deletion on chromosome 20 was mosaic in the parental unedited iPSC line (90% of cells) (see A), and two derived clones (unedited and edited) inherited it. Similarly in the experiment done at Mayo, multiple mosaic CNA in the parental line (like duplication on chromosome 1 and deletion of chromosome 14) were inherited by only one out of three clones. Additionally, one edited line, CP-601-40, was genomically unstable. In the experiment by OMRF/BCM, a mosaic duplication on chromosome 20 inherited from the unedited parental iPSC line was present in only one of the clones. Clones were selected to avoid mosaic duplication on chromosome 1 present in the parental line. **C)** Allele frequency and count of private SNVs in each clone. Clones have hundreds and thousands of point mutations that arose prior to cloning during culturing of parental lines (see Fig S4 for variant calling).

The experiment performed at the Mayo Clinic used a normal human astrocyte cell line, AA7, expressing the E6 and E7 subunits of the HPV. The line was edited to replace a fragment of DNA on chromosome 8 with a designed template sequence consisting of the same DNA fragment introducing SNP rs55705857 and a hygromycin resistance gene (for selection of cells with successful editing) (**Figs. 1A** and **S1**). Multiple clones were expanded. Upon checking the target allele with Sanger sequencing, one control (CP-581-14) and two edited clones (CP-588-36 and CP-601-40) were picked for WGS.

The OMRF/BCM experiment used an iPSC line derived from fibroblasts of a 9-year-old female patient, line c7, carrying a constitutional heterozygous variant (chr1:1,464,679 C>T; GRCh37) in exon 15 of the *ATPase family, AAA domain containing 3A* (*ATAD3A*) gene. The parental line was edited by introducing a double stranded break at the variant’s allele to correct the variant by homology directed repair. Two unedited control clones, clone7 and clone8, and one edited clone, SC20, were selected for WGS. In every experiment, we also performed WGS of the primary unedited lines.

### 2.2. Off-target effects

In each experiment, we determined off-target sites across the genome of edited lines considering all genomic sites which had up to 4 mismatched bases with respect to gRNAs. Only in the experiment performed by the Mayo Clinic, we identified two possible off-target mutations in close proximity to those off-target sites in picked clones (see **Methods**). Particularly, in the experiment by OMRF/BCM, there were two genomic sites homologous to the targeted site, represented by the paralog genes *ATAD3B* and *ATAD3C;* but no off-target editing was detected. Additionally, in the experiment by Mayo Clinic, we searched for reads supporting an insertion of the designed sequence in off-target sites and found no significant support for such an event(s) in any of the analyzed clones. We, therefore, concluded that likely no off-target editing occurred in any of the experiments.

### 2.3. Copy number alterations

In the experiment carried out at Yale, we discovered only one CNA discordant between clones (**Fig. 1B**). That was a 70 kbp heterozygous deletion in clones B10_CTR and A4 and a normal copy number region with two different haplotypes in clone A8 (**Fig. S2**). Analysis of the parental iPSC line, S1123-01-i3, revealed that the heterozygous deletion was already present in it at ~90% cell frequency. From these observations we inferred that the CNAs was mosaic in the parental, unedited line and that clones B10_CTR and A4 arose from cells with the deletion, while clone A8 from a cell without the deletion. This example demonstrates that, independently of the Cas9-induced edits, clonal lines are not necessarily isogenic due to genomic heterogeneity of cells in the parental line.

The same conclusion can be drawn from the other two experiments. In the Mayo experiment, a mosaic duplication on chromosome 1 was present in the parental line AA7 in about 10% of cells, was inherited by one of the edited clones (CP-588-36) but not by other clones. Similarly, a mosaic deletion of chromosome 14 in the parental line, AA7, was only transmitted to one of the edited clones, CP-601-40 (**Figs. 1B** and **S3**). These CNAs were totally independent from the intended Cas9-induced edits. All samples from this experiment have duplication on chromosome 20 and q arm of chromosome X, as they were constitutional to the parental line.

In the OMRF/BCM experiment, clones were selected to avoid a mosaic duplication on chromosome 1 in the parental line. But like in the above-described experiments, a mosaic duplication on chromosome 20 present in the unedited line was transmitted only to clone SC20. Again, this was independent of the intended Cas9-induced edit in *ATAD3A* gene.

### 2.4. Point mutations in individual clones

Genomic heterogeneity of parental cell line could also manifest itself in differences between clones in other variant types, so we investigated unique SNVs in each clone. To discover unique SNVs we compared clones to each other and to the parental line. In each clone SNVs inherited from the founder cell must typically have a variant allele frequency (VAF) of 50%, as the majority are present on one haplotype of autosomes. Indeed, the VAF distributions of unique SNVs in each clone were bell-shaped and centered at 50% (**Fig. 1C**), indicating that most of them represent mutations from the founder cell of the clone. Applying a cut off of 30% VAF to select SNVs likely inherited from founder (see **Methods**), we detected hundreds to thousands of SNVs in each clone (**Figs. 1C** and **S4**). Compared to iPSC clones from both Yale and OMRF/BCM cohort, the astrocyte lines (samples from Mayo), had a 10-fold higher number of SNVs, likely reflecting that the parental astrocyte line is genomically unstable. This was also reflected by the fact that clone CP-601-40 had a high fraction (~16.5%) of its genome affected by CNAs, which was outstanding even when compared to other clones of the same line. More SNVs in iPSC lines of the OMRF/BCM experiment as compared to iPSC lines of the Yale experiment may reflect earlier divergence of clones in the former case, because control clones were selected before editing and associated culture.

## 3. Discussion/Conclusions

Our study demonstrates that culture clones selected after DNA editing experiments are not isogenic and, consequently, may diverge phenotypically, independently from the intended edits. Such differences among clones arise as a consequence of the edited lines being mosaic, i.e., harboring genomic variants accumulated *in vitro.* Mosaicism in iPSC lines naturally occurs as a result of culture and could potentially be influenced by the genetic background or culture conditions favoring genomic instability (International Stem Cell et al., 2011). It was estimated that during culture, individual iPSC, intestinal, and liver stem cell lines accumulate, respectively, 3.5 ± 0.5, 7.2 ± 1.1 and 8.3 ± 3.6 base substitutions per population doubling (Kuijk et al., 2020; Thompson et al., 2020). By the time a typical experiment selects a line to be used in a CRISPR/Cas study, hundreds of point mutations are present in each cell of the line, which will be manifested, upon cloning the line, as genomic difference between clones. Most notably, clones are different in CNAs that were several kb to several mb in size. This represents, by far, the largest source of genomic divergence among clones compared to SNVs variations, as exemplified by the duplications on chromosome 1 in the OMRF/BCM and Mayo experiments. In fact, such CNAs could be comparable or even larger than the total number of genomic bases different between two random individuals or lines derived from them. Thus, clonal selection during DNA editing experiment is a large and under-appreciated source of genomic variability.

We suggest that for a meaningful comparison between edited lines, each derived clone should be subjected to WGS to screen for unintended, unique genomic variations that can have potential functional consequences. At a minimum, each clone should be screened for large CNAs to ascertain the largest source of genomic divergence between clonal lines. The functional consequences of variants outside coding regions are particularly hard to predict and could confound the interpretation of genome editing experiments. To mitigate this issue, we also propose that genome editing experiments that involve derivation of clones compare a mix of multiples unedited clones with that of edited clones.

## 4. Methods

### Experiment performed at Yale University

Multiple induced pluripotent stem cell (iPSC) lines were generated from skin fibroblasts of a 52-year-old male, S1123-01. IPSC line #3 (S1123-01-i3 at passage number 11) was selected to generate a loss of function (LOF) mutation in the *FOXG1* gene by CRISPR-Cas9 genome editing. Briefly, 4 different gRNAs targeting the beginning of the first and only exon of the *FOXG1* gene were tested for their abilities to mediate cleavage and indel formation, and the gRNA 5’ CAGGCTGTTGATGCTGAACG 3’ followed by the PAM seq 5’ AGG 3’ was chosen. Cas9/gRNA mediated 1-bp insertion was achieved by double stranded break (DSB) of DNA by Cas9 at the targeted site and error-prone repair of the break by NHEJ. Post-editing, multiple single cells were cloned and expanded. Upon genotyping at the target site by Sanger sequencing, one control (B10_CTR) and two edited clones (A4 and A8) were picked for whole genome sequencing (WGS). Selected edited clones had a 1-bp insertion (dT) into the *FOXG1* exon (at chr14:29,236,536 GTT->GTTT in GRCh37 reference genome coordinates). The insertion, heterozygous in A4 and homozygous in A8, resulted in a frameshift and premature stop codon, giving rise to a FOXG1 truncated protein. The passage numbers were 19, 19 and 21 for B10_CTR, A4 and A8 respectively. All iPSC lines were grown in mTESR1 media (StemCell Technologies) on Matrigel coated dishes (Corning Matrigel Matrix Basement Membrane Growth Factor Reduced) and propagate using Dispase (StemCell Technologies).

### Experiment performed at the Mayo Clinic

The AA7 (parental) line was subcloned from Normal Human Astrocyte (NHA) cell line expressing the E6 and E7 subunits of the HPV virus (Ohba *et al.*, 2014). Cells were grown in Dulbecco’s Modified Eagle Medium, with glutamax, 10% fetal bovine serum, and 1% Penicillin/Streptomycin. Both haplotypes in parental line AA7 have an A allele at chr8:130,645,692 (GRCh37). The line was edited with the use of a CRISPR nickases and the experiment relied on the homology directed repair to incorporate a G allele at this position. Two individual transfections were done using two different pair of gRNAs (NC1 and NC2) and the same donor sequence (**Figs. 1A**). The donor DNA contained the sequence of the regions with altered allele and the sequence of a hygromycin resistance gene flanked by LoxP sequences for removal after editing (**Fig. S1**). After transfections, cells were selected with antibiotic, subcloned and sequenced by Sanger sequencing to determine the incorporation of the G allele in the desired location. Two clones that were homozygous for the G allele at the location of interest were selected, as well as a clone that remained homozygous for A at the location of interest. All the clones that were selected were within two passages of each other at the time of editing.

### Experiment performed at OMRF/BCM

We obtained skin fibroblasts carrying the heterozygous constitutional variant (NM_001170535.1; c.1582C>T: p.Arg528Trp) in the *ATAD3A* gene from a child in a family (Harel *et al.*, 2016). Using Sendai virus-mediated reprogramming, we derived three iPSC lines (iPSC #5, #7, and #10) from the patient fibroblasts. Karyotyping G-banding showed normal genomic integrity for iPSC #7 and #10, and iPSC #7 (c7 in **Figure 1**) was selected for further studies. Two unedited control clones (clone7 and clone8) from the c7 line were selected and sequenced. The iPSC#7 was subjected to editing to correct the pathogenic variant allele (c.1582C>T) in the *ATAD3A* gene by employing CRISPR/Cas9-mediated site-specific double stranded break. We used nucleofection of ribonucleoprotein (RNP) method consisting of sgRNAs and SpCas9 protein (Liang et al., 2015) instead of vector-mediated Cas9 method. The variant specific 5’ GACGGAGGGCATGTCGGGCT 3’ gRNA and GGG PAM sequence were used. One clone SC20 with corrected pathogenic allele was picked for the analysis. The passage numbers were 21, 16, 16 and 22 for c7, clone7, clone8, and SC20 respectively.

### Sequencing data processing

Reads were aligned to the GRCh37 reference genome using BWA-Mem2 (Md et al., 2019). As a results, for each sample an alignment file in .bam format was generated. GATK-HaplotypeCaller was used to call for germline mutations.

### Checking for off-target edits

The Cas-OFFinder (Bae et al., 2014) was used to predict off-target sites using experiment specific gRNA and PAM sequence. We considered off-target site of up to four mismatch bases to each gRNA-PAM sequence. For the experiment performed at Yale, we investigated 288 possible off-target sites and found no mutations around them. For experiment performed at Mayo Clinic, two pairs of nickase gRNAs, NC1.A/NC1.B and NC2.A/NC2.B, were used (**Fig. S1**). Out of a total 1229 possible off-target sites for all nickases, only two with four mismatches had nearby mutations. The chr14:51726108:A>T mutations was nearby the genomic site similar to NC1.B gRNA (AAAaGagTTTCTTcAACAAATGG; mismatched bases are in lowercase). Similarly, mutation chr7:9805446:C>T was nearby the site similar to NC2.B gRNA (ACGGCTTCTGAtaAAaTCtTAGG; mismatched bases are in lowercase). For experiment performed at OMRF/BCM, 94 possible off-target sites were predicted and none of them had mutations.

### Checking for off-target insertion of template DNA

For the Mayo Clinic experiment, reads for each clone were re-aligned after adding a new contig to the GRCh37 reference genome. The contig contained the 150 bps of upstream homology arm, the sequence of a hygromycin resistance gene flanked by LoxP sequences, and 150 bps of downstream homology arm. We checked the reads that were aligned to the new contig and had a mate-pair read aligned to other chromosomes/contigs. As expected, most mates of the reads at the flanking region of contig mapped to chromosome 8 near the gRNA sites. Overall, 98.6% to 100% (AA7: 126/127, CP-581-14: 634/638, CP-588-36: 297/297, and CP-601-40: 378/383) of reads matched to the new contig and chromosome 8. The rest of the reads (AA7: 1/127, CP-581-14: 4/638 and CP-601-40: 5/383) aligned randomly across the genome and did not form any cluster. Hence, we concluded that no significant evidence of incorrect insertion was found. Additionally, the reads aligned to the flanking regions of the donor DNA template were checked and we found no evidence that the whole DNA template was inserted somewhere else in the genome.

### Copy number analysis

CNVpytor was used to check for copy number alterations (Suvakov et al., 2021). The mean-shift caller was used, and the read depth information was collected from the alignment files. Samples were processed with multiple bin sizes (10k, 100k). The commands were as follows:

~~~
cnvpytor -root file.pytor -rd [sample bam]
cnvpytor -root file.pytor -his 10000 100000
cnvpytor -root file.pytor -partition 10000 100000
cnvpytor -root file.pytor -call 10000 100000
cnvpytor -root file.pytor -snp file.vcf -sample sample_name
cnvpytor -root file.pytor -mask_snps
cnvpytor -root file.pytor -baf 10000 100000
~~~

We extracted read depth from alignment file for bins of 10 kbps and 100 kbps. After GC correction was performed, we used the mean-shift approach to segment read depth signal. Furthermore, we used B-allele frequency (BAF) of single nucleotide variations within called region to confirm existence of copy number alterations. Selected regions with high confidence difference from other regions are called. For each segment two p-values were calculated reflecting significance of coverage deviation from an average and deviation of BAF from 0.5. Regions with at least one of p-values smaller than 10^-20^ and with coverage difference by at least 25% from average were analyzed.

### Calling somatic point mutations

GATK-Mutect2 (Cibulskis et al., 2013) was used to call for somatic mutations. The following command was used:

~~~
GATK4 --java-options “-Xmx12G -Djava.io.tmpdir=$OUTDIR/tmp -XX:-UseParallelGC” Mutect2 \
        -R $REF_GENOME -I $TUMOR_BAM -I $NORMAL_BAM -normal $NORMAL_SAMPLE \
        -O $OUTDIR/${NORMAL_SAMPLE}_${TUMOR_SAMPLE}.mutect.vcf.gz
GATK4 --java-options “-Xmx12G -Djava.io.tmpdir=$OUTDIR/tmp -XX:-UseParallelGC” \ FilterMutectCalls -R $REF_GENOME \
        -V $OUTDIR/${NORMAL_SAMPLE}_${TUMOR_SAMPLE}.mutect.vcf.gz \
        -O $OUTDIR/${NORMAL_SAMPLE}_${TUMOR_SAMPLE}.mutect.filtered.vcf.gz
~~~

GATK version: 4.1.8.1; Reference Genome: GRCh37

In each clone, mutations were called by comparing against other clones and against bulk sample from the parental line. We then considered the overlapping set of calls from the three comparisons as culture-derived somatic mutations in the parental line transmitted to each of the clones (**Fig. S4**). Such mutations must have VAF distribution looking like a symmetrical bell-shaped curve centered around 50%, i.e., almost all mutations are on one haplotype of autosomes. Indeed, we observed such a shape (**Fig. 1C**). To eliminate mutation that may have arisen during culturing of clones, we selected only mutations with at least 30% VAF.

### Ethics statement

For experiments performed at Yale University, samples were collected from de-identified individuals. The research study was approved by the Yale University Institutional Review Board. The BioBank protocols are in accordance with the ethical standards of Yale University. For experiments performed at OMRF/BCF, the family provided consent according to the Baylor-Hopkins Center for Mendelian Genomics (BHCMG) research protocol H-29697, approved by the Institutional Review Board (IRB) at Baylor College of Medicine (BCM).

## 5. Data Availability

Sequencing data is being deposited to DbGaP and will become available for qualified researchers as soon as the processing is complete.

## Notes

### Competing Interest Statement

The authors have declared no competing interest.

## References

Assou, S., Bouckenheimer, J., and De Vos, J. (2018). Concise Review: Assessing the Genome Integrity of Human Induced Pluripotent Stem Cells: What Quality Control Metrics? Stem Cells 36, 814–821. 10.1002/stem.2797.

Bae, S., Park, J., and Kim, J.-S. (2014). Cas-OFFinder: a fast and versatile algorithm that searches for potential off-target sites of Cas9 RNA-guided endonucleases. Bioinformatics 30, 1473–1475. 10.1093/bioinformatics/btu048.

Boyle, E.A., Andreasson, J.O.L., Chircus, L.M., Sternberg, S.H., Wu, M.J., Guegler, C.K., Doudna, J.A., and Greenleaf, W.J. (2017). High-throughput biochemical profiling reveals sequence determinants of dCas9 off-target binding and unbinding. Proc Natl Acad Sci U S A 114, 5461–5466. 10.1073/pnas.1700557114.

Cibulskis, K., Lawrence, M.S., Carter, S.L., Sivachenko, A., Jaffe, D., Sougnez, C., Gabriel, S., Meyerson, M., Lander, E.S., and Getz, G. (2013). Sensitive detection of somatic point mutations in impure and heterogeneous cancer samples. Nat Biotechnol 31, 213–219. 10.1038/nbt.2514.

Fu, Y., Foden, J.A., Khayter, C., Maeder, M.L., Reyon, D., Joung, J.K., and Sander, J.D. (2013). High-frequency off-target mutagenesis induced by CRISPR-Cas nucleases in human cells. Nature Biotechnology 31, 822–826. 10.1038/nbt.2623.

Gaj, T., Gersbach, C.A., and Barbas, C.F., 3rd (2013). ZFN, TALEN, and CRISPR/Cas-based methods for genome engineering. Trends Biotechnol 31, 397–405. 10.1016/j.tibtech.2013.04.004.

Hanna, R.E., and Doench, J.G. (2020). Design and analysis of CRISPR-Cas experiments. Nat Biotechnol 38, 813–823. 10.1038/s41587-020-0490-7.

Harel, T., Yoon, W.H., Garone, C., Gu, S., Coban-Akdemir, Z., Eldomery, M.K., Posey, J.E., Jhangiani, S.N., Rosenfeld, J.A., Cho, M.T., et al. (2016). Recurrent De Novo and Biallelic Variation of ATAD3A, Encoding a Mitochondrial Membrane Protein, Results in Distinct Neurological Syndromes. Am J Hum Genet 99, 831–845. 10.1016/j.ajhg.2016.08.007.

International Stem Cell, I., Amps, K., Andrews, P.W., Anyfantis, G., Armstrong, L., Avery, S., Baharvand, H., Baker, J., Baker, D., Munoz, M.B., et al. (2011). Screening ethnically diverse human embryonic stem cells identifies a chromosome 20 minimal amplicon conferring growth advantage. Nat Biotechnol 29, 1132–1144. 10.1038/nbt.2051.

Jinek, M., Chylinski, K., Fonfara, I., Hauer, M., Doudna, J.A., and Charpentier, E. (2012). A programmable dual-RNA-guided DNA endonuclease in adaptive bacterial immunity. Science 337, 816–821. 10.1126/science.1225829.

Kim, D., Bae, S., Park, J., Kim, E., Kim, S., Yu, H.R., Hwang, J., Kim, J.I., and Kim, J.S. (2015). Digenome-seq: genome-wide profiling of CRISPR-Cas9 off-target effects in human cells. Nat Methods 12, 237–243, 231 p following 243. 10.1038/nmeth.3284.

Kim, Y.G., Cha, J., and Chandrasegaran, S. (1996). Hybrid restriction enzymes: zinc finger fusions to Fok I cleavage domain. Proc Natl Acad Sci U S A 93, 1156–1160. 10.1073/pnas.93.3.1156.

Kuijk, E., Jager, M., van der Roest, B., Locati, M.D., Van Hoeck, A., Korzelius, J., Janssen, R., Besselink, N., Boymans, S., van Boxtel, R., and Cuppen, E. (2020). The mutational impact of culturing human pluripotent and adult stem cells. Nat Commun 11, 2493. 10.1038/s41467-020-16323-4.

Liang, X., Potter, J., Kumar, S., Zou, Y., Quintanilla, R., Sridharan, M., Carte, J., Chen, W., Roark, N., Ranganathan, S., et al. (2015). Rapid and highly efficient mammalian cell engineering via Cas9 protein transfection. J Biotechnol 208, 44–53. 10.1016/j.jbiotec.2015.04.024.

Ma, E., Harrington, L.B., O’Connell, M.R., Zhou, K., and Doudna, J.A. (2015). Single-Stranded DNA Cleavage by Divergent CRISPR-Cas9 Enzymes. Mol Cell 60, 398–407. 10.1016/j.molcel.2015.10.030.

Md, V., Misra, S., Li, H., and Aluru, S. (2019). Efficient Architecture-Aware Acceleration of BWA-MEM for Multicore Systems. 2019 Ieee Int Parallel Distributed Process Symposium Ipdps 00, 314–324. 10.1109/ipdps.2019.00041.

Ohba, S., Mukherjee, J., See, W.L., and Pieper, R.O. (2014). Mutant IDH1-driven cellular transformation increases RAD51-mediated homologous recombination and temozolomide resistance. Cancer Res 74, 4836–4844. 10.1158/0008-5472.CAN-14-0924.

Pattanayak, V., Lin, S., Guilinger, J.P., Ma, E., Doudna, J.A., and Liu, D.R. (2013). High-throughput profiling of off-target DNA cleavage reveals RNA-programmed Cas9 nuclease specificity. Nat Biotechnol 31, 839–843. 10.1038/nbt.2673.

Ran, F.A., Hsu, P.D., Wright, J., Agarwala, V., Scott, D.A., and Zhang, F. (2013). Genome engineering using the CRISPR-Cas9 system. Nat Protoc 8, 2281–2308. 10.1038/nprot.2013.143.

Suvakov, M., Panda, A., Diesh, C., Holmes, I., and Abyzov, A. (2021). CNVpytor: a tool for copy number variation detection and analysis from read depth and allele imbalance in whole-genome sequencing. GigaScience 10. 10.1093/gigascience/giab074.

Thompson, O., von Meyenn, F., Hewitt, Z., Alexander, J., Wood, A., Weightman, R., Gregory, S., Krueger, F., Andrews, S., Barbaric, I., et al. (2020). Low rates of mutation in clinical grade human pluripotent stem cells under different culture conditions. Nat Commun 11, 1528. 10.1038/s41467-020-15271-3.

Tsai, S.Q., Nguyen, N.T., Malagon-Lopez, J., Topkar, V.V., Aryee, M.J., and Joung, J.K. (2017). CIRCLE-seq: a highly sensitive in vitro screen for genome-wide CRISPR-Cas9 nuclease off-targets. Nat Methods 14, 607–614. 10.1038/nmeth.4278.

