## Supplementary Data for "Clonally selected lines after CRISPR/Cas editing are not isogenic"

**Supplementary Figure S1**: The CRISPR-Cas9 design for the experiment performed at Mayo Clinic. Two pairs of gRNAs (NC1 and NC2) were used to introduce single stranded break on four sites. Individual transfections were done using either one of the pairs, and clones were picked, genotyped for incorporation of the template DNA and sequenced. Two clones, CP-581-14 and CP-588-36 were picked from transfection with NC1 and one clone, CP-601-40, was picked from transfection with NC2. The gRNAs for nickase construct 1 (NC1) target two positions NC1.A (chr8:130, 645,666, reverse strand; GRCh37) and NC1.B (chr8:130, 545, 712, forward strand; GRCh37). Similarly, the gRNAs for nickase construct 2 (NC2) target position NC2.A (chr8: 130, 645,685; reverse strand; GRCh37) and NC2.B (chr8:130, 545, 728, forward strand; GRCh37) on each strand. The breaks were repaired by homology directed repair using donor sequence to incorporate a SNV (chr8:130,645,692 A>G; GRCh37). The donor sequence includes homology sequence with an alternate allele and the hygromycin gene flanked by LoxP sequences.

­_­­­­­_

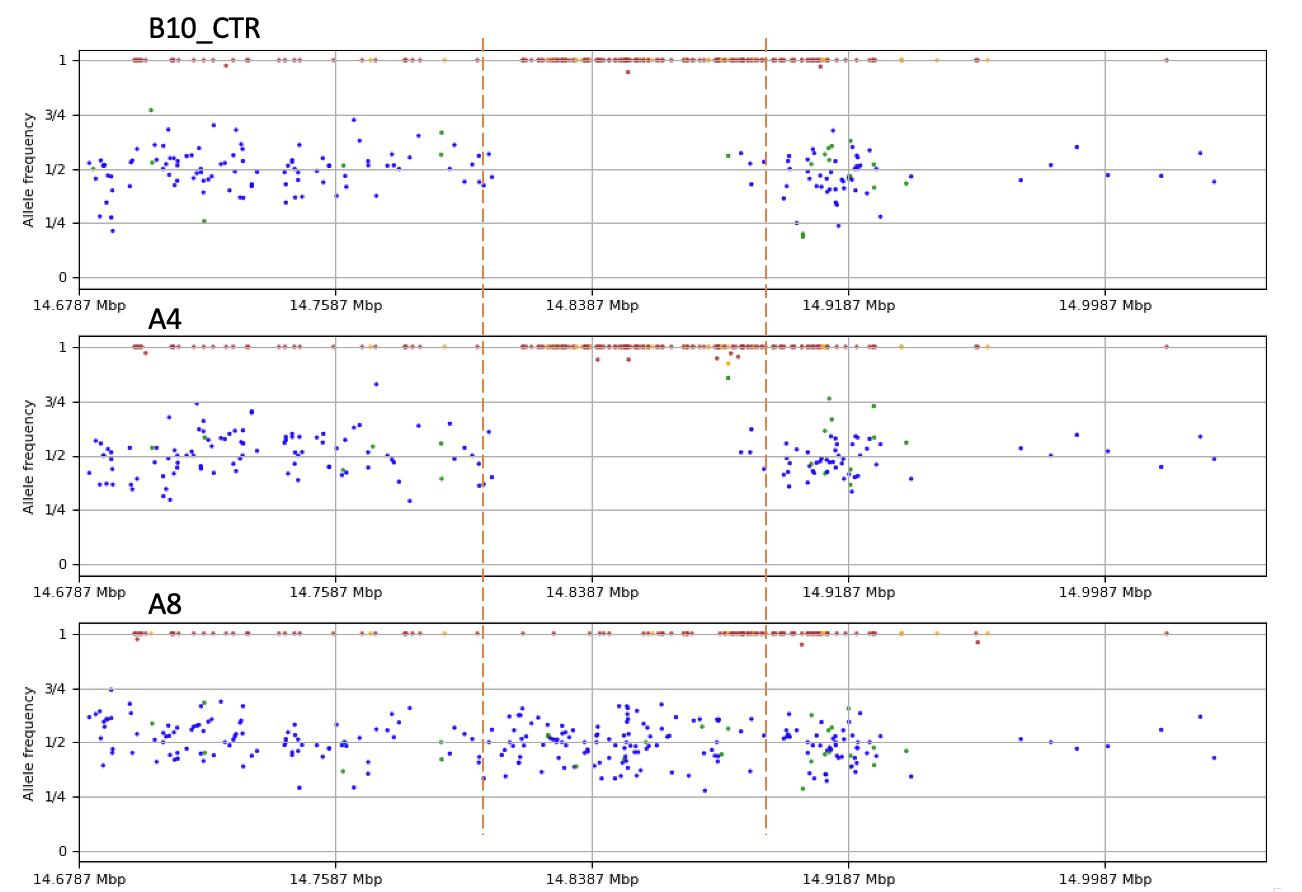

**Supplementary Figure S2**: Details for the copy number altered region chr20:14678706-15040811 (dashed vertical lines) for derived clones in the experiment by Yale. A8 has heterozygous SNPs (blue dots) within the regions, indicating presence of two different haplotypes. Other clones, B10_CTR and A4, have only homozygous SNPs (red dots). Read depth profile indicates that these two clones have heterozygous deletion (see **Fig. 1**).

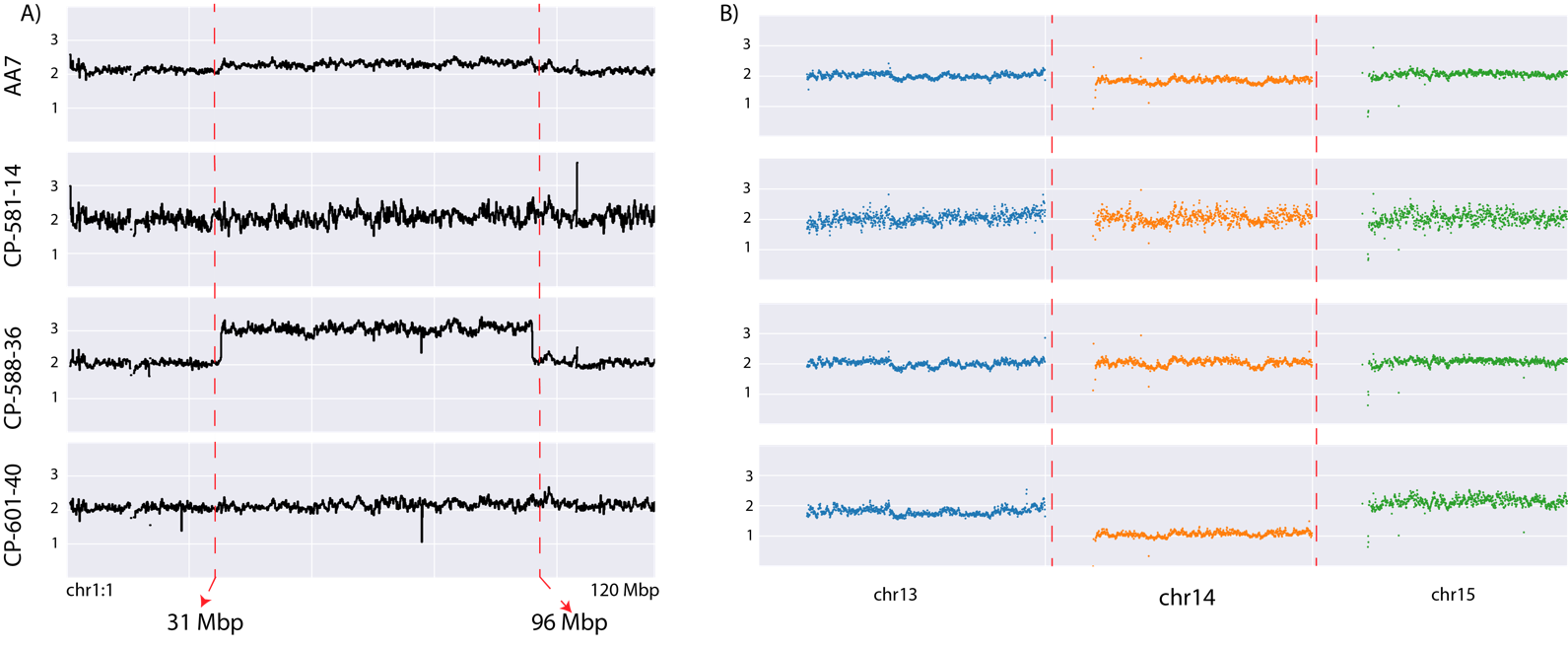
**Supplementary Figure S3:** Parental mosaic CNAs were inherited in one of three subclones for experiments performed at the Mayo Clinic (y-axis is estimated copy number). **A)** About 10% cells of unedited parental line, AA7, had a duplication at chr1:31mb-96mb which was inherited by the CP-588-36 clone**. B)** About 10% cells of unedited parental line AA7 had a deletion on chromosome 14, which was inherited by the CP-601-40 clone.

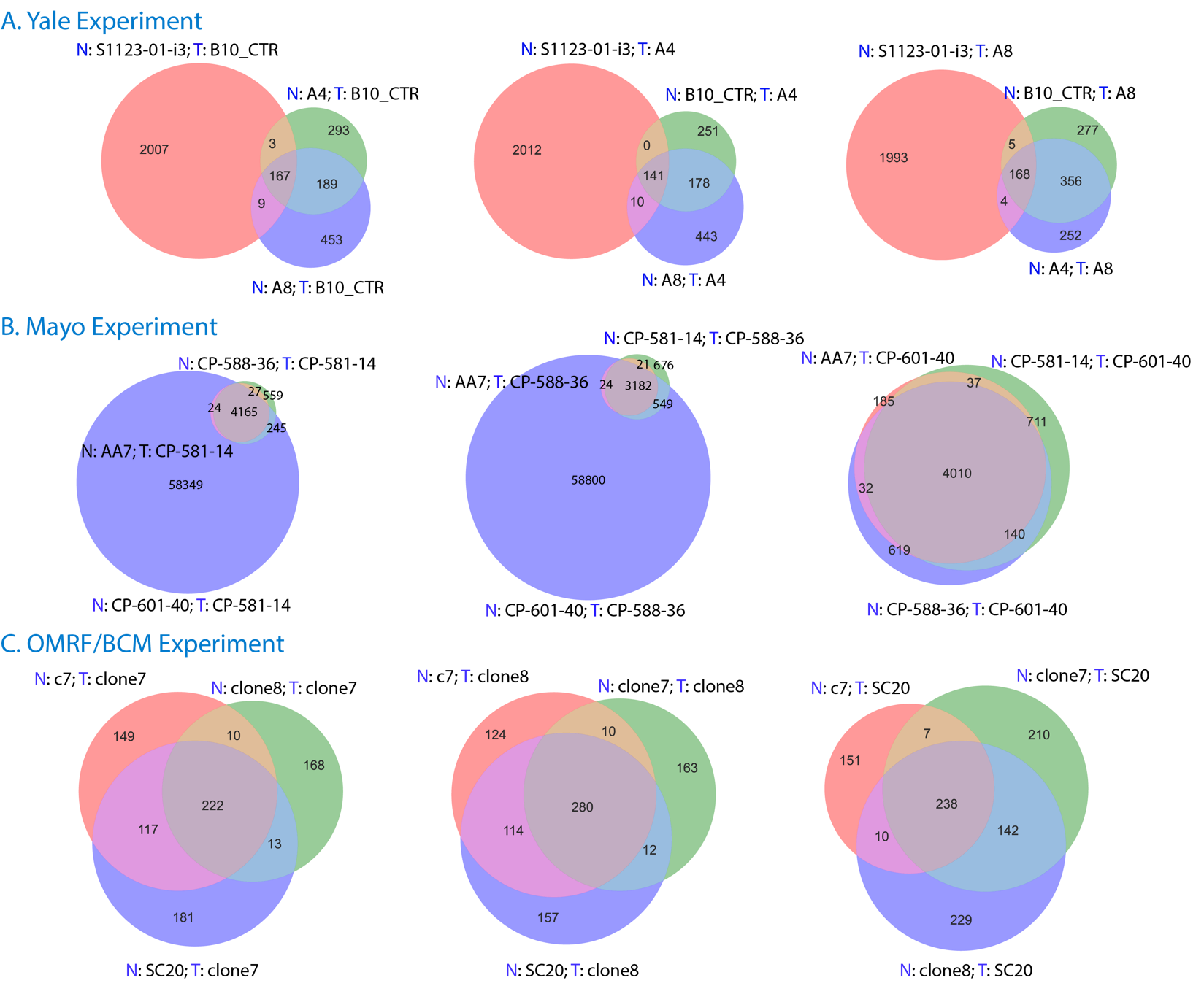

**Supplementary Figure S4**: Unique single nucleotide mutations (SNVs) with variant allele frequency above 30% for each derived clone. Each Venn diagram corresponds to SNVs calls in one clone. Each circle in the diagram represents a comparison between a clone (named after T:) and other clone/sample (named after N:). For each clone unique mutations were taken as calls from the intersection of three comparisons. **A)** For B10_CTR, A4, and A8 clones 167, 141 and 168 unique mutations were called. **B)** CP-581-14, CP-588-35 and CP-601-40 clones had 4165, 3182 and 4010 unique mutations. **C**) 238, 222 and 280 unique mutations were called for SC20, clone7 and clone8 clones respectively.
